# From the shallows to the depths: A new probe set to target ultraconserved elements for Malacostraca

**DOI:** 10.1101/2023.06.30.547307

**Authors:** Jonas C. Geburzi, Paula C. Rodríguez-Flores, Shahan Derkarabetian, Gonzalo Giribet

## Abstract

Since its introduction about a decade ago, target enrichment sequencing of ultraconserved elements (UCEs) has proven to be an invaluable tool for studies across evolutionary scales, and thus employed from population genetics, to historical biogeography and deep phylogenetics. UCE probe sets are available for an increasing range of major taxonomic groups, including cnidarians, vertebrates, terrestrial arthropods, and mollusks. Here, we present the first probe set targeting UCEs in crustaceans, specifically designed for decapods and other malacostracan lineages. Probes were designed using published genomes of nine decapod and one peracarid species, as well as raw Nanopore long reads of one additional brachyuran species. The final probe set consists of about 20,000 probes, targeting 1,348 unique UCE loci. Preliminary analyses of UCE data obtained from an intertidal mangrove crab, and from deep-sea squat lobsters indicate high UCE recovery rates (about 1,000 loci per sample) in evolutionarily shallow datasets. To test the probe set at deeper phylogenetic levels, we compiled a dataset across Malacostraca (including representatives of Decapoda, Peracarida, Euphausiacea, Stomatopoda, and Phyllocarida), and were able to recover hundreds of UCEs for the non-decapod species, expanding the targeted use of this UCE probeset to all Malacostraca. Additionally, we recovered similar numbers of UCEs from historical museum specimens up to > 150 years old, that were included in all datasets, confirming that UCEs are a fruitful technique for leveraging museum specimens for genomic studies. Overall, our results highlight the versatility of this UCE probe set and its high potential for crustacean evolutionary studies.

## 1 Introduction

Ultraconserved elements (UCEs) are highly conserved genome regions that are present across many taxa, and are known to have regulatory expression functions in vertebrates, while being of exonic origin in arthropods (Faircloth et al., 2012; Hedin et al., 2019; McCormack et al., 2012). Since their discovery as syntenically conserved regions in the human, mouse and rat genomes (Bejerano et al., 2004), the use of UCEs for phylogenomic studies has rapidly increased, becoming a popular technique in the past few years (see references below). The increasing popularity of the UCEs approach relies on 1) the amount of generated data, since it allows to target between hundreds to thousands of orthologous loci, improving phylogenetic resolution and largely outperforming traditional multilocus sequencing; 2) in contrast to other genomic techniques such as transcriptomics, UCE sequencing allows to obtain historical DNA from ethanol preserved specimens in museum collections (Blaimer et al., 2016; Derkarabetian et al., 2019; McCormack et al., 2016; Raxworthy & Smith, 2021); and, 3) unlike Anchored Hybrid Enrichment (AHE; Lemmon et al., 2012), the full UCE probe set design and hybridization pipeline was originally published open source (Faircloth, 2017), allowing everybody to design baits to target their organisms of study.

UCEs have been used to reconstruct phylogenies for many taxa across the whole Tree of Life and at multiple phylogenetic scales; from backbone phylogenies (e.g., Faircloth et al., 2013; Streicher & Wiens, 2017) to species-level or even population-level studies (e.g., Derkarabetian et al., 2018, 2022; Smith et al., 2014). The versatility of UCEs at different evolutionary scales relies on the sequencing of UCEs flanking regions, with a higher proportion of variable sites when increasing the distance from the highly conserved core of the UCE (Faircloth et al., 2012). This allows generation of conserved datasets useful for phylogenomic reconstruction at higher taxonomic levels, as well as the possibility to extract single nucleotide polymorphisms (SNPs) from each single locus, showing an efficacy comparable to microsatellites and ddRAD for population genomics (Glon et al., 2021; Vinciguerra et al., 2019).

UCE probe sets are currently available for several invertebrate taxa, and at different levels of divergence, from phylum to genus-level: e.g., for Cnidaria (Quattrini et al., 2018), Hexacorallia (Cowman et al., 2020), for Mollusca Heterobranchia (Moles & Giribet, 2021) and for multiple arthropod taxa, including several groups of insects and arachnids (Starrett et al., 2017; Zhang et al., 2019; and references therein). Within arthropods, the most comprehensive probe set was designed for Arachnida, a group spanning more than 500 million years of evolution (Starrett et al., 2017). However, despite this huge divergence, this probeset has proven to be very useful across all chelicerate groups and across variety of phylogenetic levels (e.g., Ballesteros et al., 2021; Boyer et al., 2022; de Miranda et al., 2022; Hedin et al., 2020).

A major component of Earth’s Biodiversity is represented by crustaceans of the Class Malacostraca. This group has a long evolutionary history, and like arachnids, probably started to diversify at least 500 million years ago (Bernot et al., 2022; Schram & Abele, 1982; Schwentner et al., 2017). It is the largest class of non-hexapod crustaceans with more than 30,000 described species, classified into 17 orders. Malacostracans display great morphological disparity, including multiple body forms such as shrimp-like, crab-like, lobster-like, bivalved crustaceans, mantis-like, conglobated forms, etc. This group has received much attention due to their ancient origin, morphological disparity and ecological diversity. However, although it is considered a monophyletic group, the scarcity of genomic studies has hampered the understanding of the phylogenetic placement of many malacostracan groups (e.g., Höpel et al., 2022; Schwentner et al., 2018; Shen et al., 2015). Additionally, within Malacostraca, decapods and peracarids are hyperdiverse taxa with independent radiations to the land and fresh waters (e.g., Hou et al., 2014; Tsang et al., 2022; Wolfe et al., 2023). They have colonized extreme environments in both marine and terrestrial habitats, such as anchialine caves, hydrothermal vents or hadal trenches (e.g., Gonzalez et al., 2020; Patel et al., 2020; Swan et al., 2021), and therefore represent a model group for several disciplines in evolutionary biology.

Despite the rapid growth of UCEs to study evolution of arthropod lineages, no crustacean probe set currently exists. The use of target capture sequencing of UCEs in crustacean research can offer multiple benefits at various levels. Firstly, it can aid in resolving phylogenetic uncertainties in Malacostraca, providing insights into the evolutionary relationships of this diverse group of crustaceans. Secondly, it can improve phylogenetic resolution for the study of explosive radiations and other evolutionary events at different scales, ranging from intra-specific to interspecific levels. Finally, this approach can be especially valuable in cases where access to living specimens is limited and expensive, such as with deep-sea samples. By utilizing historical collections, researchers can leverage this technique to solve taxonomic questions using molecular methods, thereby advancing our understanding of this fascinating group of organisms and adding value to the extensive museum collections of deep sea fauna and regions of the world where collecting is no longer feasible.

Here we present a new probe set targeting UCEs for Decapoda, which is the first UCE probe set for any crustacean lineage to date. We tested this probe set *in-silico* and *in-vitro* by hybridization to DNA samples obtained from all major decapod lineages as well as several non-decapod Malacostraca. Our tests demonstrate the efficacy of the probe set at different taxonomic scales, regularly recovering hundreds to more than 1,000 loci across the Malacostraca, and still dozens to hundreds of loci from historical samples (collection dates 1865 to 2012) in our test datasets. The new probe set will thus provide a valuable resource for population-level to phylogenomic studies across Malacostraca.

## 2 Material and Methods

### 2.1 Taxon sampling

The publicly available decapod genomes cover the majority of the order’s lineages. For the probe set design, we used seven genomes from across the Decapoda tree of life, with a focus on the most diverse group, the true crabs (Brachyura), our main focal taxon. The mitten crab *Eriocheir sinensis* (Brachyura, Grapsoidea) was used as base genome, *Birgus latro* (Anomura), *Chionoecetes opilio* (Brachyura, Majoidea), *Homarus americanus* (Astacidea), *Macrobrachium nipponense* (Caridea), *Panulirus ornatus* (Achelata) and *Penaeus japonicus* (Dendrobranchiata) were used as exemplar taxa. The peracarid *Hyalella azteca* (Amphipoda) was used as outgroup taxon. Additionally, we added novel Oxford Nanopore raw reads of a third crab species, *Aratus pisonii* (Brachyura, Grapsoidea) to the probe set design pipeline.

We downloaded 13 additional crustacean genomes from NCBI to test the probe set *in-silico*. These included seven decapods, four peracarids, and two non-malacostracan genomes (one branchiopod and one copepod), to also assess performance and efficacy of the probe set at deeper phylogenetic levels (Table S1).

For the *in-vitro* test of the probe set, we compiled 41 samples from the collections of the Museum of Comparative Zoology, Cambridge, MA (MCZ), the Smithsonian National Museum of Natural History, Washington DC (USNM), and the Musée national de l’Histoire naturelle, Paris (MNHN), focusing on the Decapoda, but also including representatives of Peracarida (Amphipoda, Cumacea, Isopoda), Euphausiacea, Stomatopoda, Phyllocarida, and Branchiopoda (Table 1). Ten of these samples were obtained from “historical” specimens with degraded DNA, i.e. collected more than 10 years ago, and stored in 70% ethanol at room temperature. The historical samples covered a range of collection years between 2012 and 1865, and were included to assess UCE locus recovery and utility of the probe set for phylogenetic studies including crustacean specimens from museum collections. These 41 *in-vitro* and 21 *in-silico* malacostracan samples combined are subsequently referred to as the “Malacostraca dataset”.

**Table 1.** Assembly and UCE extraction summary statistics for the *in-vitro* samples in the Malacostraca dataset, including the number of assembled contigs, the number of raw UCE loci recovered from contigs, the length of the raw UCE loci, and the final number of UCE loci in a 50% taxon coverage matrix after trimming and filtering

| Higher taxonomy | Species | Voucher | Year | contigs | UCEs (raw) | Locus length (bp) |  |  | UCEs (final) |
| --- | --- | --- | --- | --- | --- | --- | --- | --- | --- |
|  |  |  |  |  |  | mean | min | max |  |
| Brachyura |  |  |  |  |  |  |  |  |  |
| Dynomenidae | *Metadynomene* sp. | MCZ:IZ:153343 | 2019 | 83,578 | 1,166 | 611.8 | 236 | 1,157 | 847 |
| Gecarcinidae | *Gecarcinus ruricola* | MCZ:IZ:46902 | 2003 | 1,982 | 48 | 258.0 | 80 | 496 | 29 |
| Grapsidae | *Pachygrapsus transversus* | MCZ:IZ:61995 | 2004 | 38,220 | 1,069 | 455.0 | 227 | 3,619 | 741 |
| Percnidae | *Percnon planissimum* | MCZ:IZ:163892 | 2022 | 463,250 | 657 | 1,399.9 | 394 | 5,678 | 528 |
| Portunidae | *Callinectes sapidus* | MCZ:IZ:23424 | 2010 | 94,335 | 290 | 329.4 | 80 | 6,602 | 16 |
| Sesarmidae | *Aratus pisonii* | MCZ:IZ:161758 | 2021 | 239,522 | 1,031 | 1,503.0 | 77 | 10,525 | 788 |
|  | *Aratus pisonii* | MCZ:IZ:6197 | 1865 | 2552 | 5 | 212.6 | 80 | 257 | 3 |
|  | *Armases cinereum* | MCZ:IZ:162859 | 2021 | 417,860 | 744 | 1,217.5 | 245 | 5,048 | 589 |
| Trapeziidae | *Trapezia rufopunctata* | MCZ:IZ:163860 | 2022 | 108,611 | 847 | 1,622.1 | 247 | 3,786 | 656 |
| Varunidae | *Eriocheir sinensis* | MCZ:IZ:68008 | 1932 | 10,746 | 40 | 265.5 | 79 | 493 | 13 |
| Xanthidae | *Actaeodes tomentosus* | MCZ:IZ:163780 | 2022 | 794,710 | 243 | 1,146.9 | 269 | 2,115 | 198 |
|  | *Cymo melanodactylus* | MCZ:IZ:163854 | 2022 | 578,024 | 601 | 841.8 | 205 | 4,203 | 463 |
| Anomura |  |  |  |  |  |  |  |  |  |
| Eumunididae | *Eumunida* cf. *pacifica* | MCZ:IZ:153356 | 2019 | 510,382 | 1,051 | 1,255.2 | 81 | 5,316 | 793 |
|  | *Eumunida pacifica* | MNHN-IU-2014-23737 | 1991 | 110,384 | 975 | 398.7 | 229 | 6,054 | 656 |
|  | *Pseudomunida fragilis* | MCZ:IZ:151084 | 2018 | 84,402 | 965 | 1,275.2 | 229 | 6,592 | 730 |
| Galatheididae | *Coralliogalathea parva* | MCZ:IZ:163893 | 2022 | 355,095 | 612 | 1,091.5 | 240 | 6,396 | 466 |
|  | *Galathea platycheles* | MCZ:IZ:163971 | 2022 | 733,875 | 510 | 1,118.5 | 280 | 2,322 | 412 |
| Munididae | *Grimothea quadrispina* | MCZ:IZ:139215 | 2016 | 1,108,273 | 531 | 873.5 | 235 | 1,528 | 415 |
|  |  |  |  |  |  | mean | min | max |  |
| Munidopsidae | *Munidopsis lentigo* | USNM 1487884 | 2012 | 423,853 | 1000 | 1,043.2 | 232 | 8,073 | 748 |
| Paguridae | *Porcellanopagurus* sp. | MNHN-IU-2021-7149 | 2021 | 4,229 | 108 | 295.9 | 80 | 575 | 65 |
| Porcellanidae | *Eucramus transversilineatus* | MCZ:IZ:30775 | 2011 | 82,661 | 735 | 498.0 | 194 | 1,815 | 550 |
| **Axiidea** |  |  |  |  |  |  |  |  |  |
| Axiidae | Axiidae sp. 1 | MCZ:IZ:150640 | 2016 | 138,693 | 957 | 650.3 | 229 | 4,892 | 664 |
|  | Axiidae sp. 2 | MCZ:IZ:163883 | 2022 | 157,219 | 837 | 1,083.0 | 229 | 13,111 | 623 |
| **Astacidea** |  |  |  |  |  |  |  |  |  |
| Cambaridae | *Procambarus clarkii* | MCZ:IZ:68053 | 2015 | 188,529 | 913 | 819.6 | 276 | 5,449 | 675 |
| Nephropidae | *Homarus americanus* | MCZ-IZ162632 | 2019 | 55,790 | 918 | 1,672.0 | 323 | 8,079 | 711 |
| **Polychelida** |  |  |  |  |  |  |  |  |  |
| Polychelidae | *Pentacheles laevis* | MCZ:IZ:19151 | 2012 | 181,358 | 669 | 1,161.5 | 232 | 6,871 | 528 |
| **Caridea** |  |  |  |  |  |  |  |  |  |
| Alpheidae | *Synalpheus* sp. | MCZ:IZ:163842 | 2022 | 134,448 | 1,131 | 679.3 | 229 | 1,712 | 797 |
| Palaemonidae | *Zenopontonia soror* | MCZ:IZ:163834 | 2022 | 98,044 | 1,018 | 1,131.9 | 231 | 4,138 | 768 |
| **Stenopodidea** |  |  |  |  |  |  |  |  |  |
| Stenopodidae | *Stenopus hispidus* | MCZ:IZ:163857 | 2022 | 417,365 | 842 | 890.0 | 186 | 1,933 | 602 |
| **Dendrobranchiata** |  |  |  |  |  |  |  |  |  |
| Penaeidae | *Penaeus japonicus* | MCZ:IZ:9630 | 1933 | 4,698 | 38 | 251.4 | 80 | 393 | 3 |
| Sergestidae | *Robustosergia robusta* | MCZ:IZ:46338 | 2014 | 541,292 | 441 | 1,296.3 | 230 | 4,106 | 349 |
| **Euphausiacea** |  |  |  |  |  |  |  |  |  |
| Euphausiidae | *Euphausia pacifica* | MCZ:IZ:148541 | 2018 | 86,011 | 828 | 1,034.8 | 157 | 7,162 | 573 |
| **Stomatopoda** |  |  |  |  |  |  |  |  |  |
| Gonodactylidae | *Gonodactylellus* sp. | MCZ:IZ:163794 | 2022 | 311,979 | 906 | 1,052.6 | 230 | 5,658 | 658 |
| **Isopoda** |  |  |  |  |  |  |  |  |  |
| Armadillidiidae | *Armadillidium vulgare* | MCZ:IZ:28779 | 2007 | 18,254 | 204 | 293.4 | 80 | 2,604 | 31 |
|  |  |  |  |  |  | mean | min | max |  |
| Idoteidae | *Idotea balthica* | MCZ:IZ:150668 | 2018 | 154,544 | 799 | 1,175.0 | 235 | 10,191 | 612 |
| **Amphipoda** |  |  |  |  |  |  |  |  |  |
| Hyalellidae | *Hyalella azteca* | MCZ:IZ:78322 | 1969 | 2,767 | 123 | 276.4 | 80 | 1,039 | 3 |
| Maeridae | Maeridae sp. | MCZ:IZ:163784 | 2022 | 278,164 | 820 | 986.1 | 140 | 2,381 | 571 |
| **Cumacea** | Cumacea sp. | MCZ:IZ:742248 | 2015 | 215,981 | 454 | 927.6 | 83 | 2,948 | 285 |
| **Lophogastrida** |  |  |  |  |  |  |  |  |  |
| Gnathophausiidae | *Gnathophausia zoea* | MCZ:IZ:46341 | 2014 | 41,250 | 500 | 324.0 | 80 | 911 | 348 |
| **Leptostraca** |  |  |  |  |  |  |  |  |  |
| Nebaliidae | *Nebalia* sp. | MCZ:IZ:126457 | 2004 | 5,896 | 72 | 278.2 | 79 | 586 | 14 |
| **Anostraca** |  |  |  |  |  |  |  |  |  |
| Artemiidae | *Artemia franciscana* | MCZ:IZ:58866 | 2001 | 32,021 | 196 | 283.9 | 79 | 487 | 111 |
| **mean fresh** |  |  |  | **307,115** | **762** | **1,040.7** | **212** | **4,926** | **568** |
| **mean total** |  |  |  | **227,094** | **632** | **828.8** | **183** | **4,081** | **454** |

Furthermore, we compiled two additional datasets to assess the efficacy of the probe set at shallower taxonomic levels, i.e. its potential to recover variation among closely related species and populations of a single species: A set of 48 samples representing 30 species of the squat lobster family Eumunididae (Anomura), collected between 1906 and 2022, including historical specimens from the MCZ, USNM, MNHN, and the California Academy of Sciences, San Francisco (CAS) (the “*Eumunida* dataset” in the following, Table S2); and 23 samples of the mangrove crab *Aratus pisonii* (Brachyura, Sesarmidae), collected between 1859 and 2021 around Miami, FL, including historical specimens from the MCZ, USNM, and the Florida Museum of Natural History, Gainesville (UF) (the “*Aratus* dataset” in the following, Table S3).

### 2.2 Probe set design and synthesis

We used PHYLUCE version 1.7.1 (Faircloth, 2016) to design the probe set, following the tutorial on https://phyluce.readthedocs.io/en/latest/tutorials/tutorial-4.html (also see Faircloth, 2017). The downloaded genomes were converted to FASTA format and file headers were modified for compatibility with the PHYLUCE pipeline using SeqIO, available on Bioconda (Grüning et al., 2018). Copies of the genomes in 2bit format, required further downstream in the pipeline, were created using faToTwoBit (Kent, 2002). ART (Huang et al., 2012) was then used to simulate 100-bp short reads with 200-bp insert size (150 standard deviation) from all exemplar genomes and the *A. pisonii* long reads, covering the base genome approximately 2X. These simulated reads were individually aligned to the base genome using STAMPY (Lunter & Goodson, 2011), identifying putatively conserved loci with less than 5% sequence divergence to the base genome. Unmapped reads were removed from the alignment files using the *view* function in SAMtools (Li et al., 2009), and the cleaned alignments were converted to BED format, sorted by position along scaffolds, and proximate contigs were merged using BEDTools (Quinlan & Hall, 2010). Finally, phyluce_probe_strip_masked_loci_from_set in PHYLUCE was used to filter out putatively conserved loci that are repetitive regions, by removing all contigs shorter than 80 bp and where more than 25% of the base genome were masked.

Based on these pairwise alignment files, an SQLite database was created to query conserved loci shared across several taxa. From this database, we identified those loci shared between the base genome and six out of seven exemplar genomes, and extracted sequences corresponding to these loci from the base genome, buffered to 160 bp, with phyluce_probe_get_genome_sequences_from_bed. For each of these sequences, two 120-bp probes with 40 bp overlap were designed to achieve 3X tiling density at the center of the conserved locus. Potentially problematic probes with more than 25% masked bases, below 30% or above 70% GC content, as well as potential duplicate probes with more than 50% coverage and identity were subsequently removed, to create a temporary probe set.

To design the final probe set, the temporary probes were aligned back to all exemplar genomes, now also including the outgroup *Hyalella azteca*, with a minimum identity of 50%. Sequences of 180 bp length were extracted from the targeted conserved loci in all taxa and written to FASTA files. Another SQLite database with the matches of the temporary probes to conserved loci across all taxa was created and used to identify loci that were recovered by a temporary probe in at least seven out of the nine taxa (7 exemplar, 1 outgroup, 1 base genome). For each of these loci, two 120-bp probes with 3X tiling density were designed from all taxa, filtered and duplicates removed as described above. *In-silico* tests of the UCE probe set were performed by assessing locus recovery across a range of decapod and non-decapod crustacean genomes, using phyluce_assembly_match_contigs_to_probes after aligning the probe set to these genomes (see Table S1).

The concatenated probe set file was sent to Arbor BioSciences, where final tests of the design were performed before synthesizing the probes. Each probe was BLASTed against the base genome, and non-specific probes, as well as probes targeting over-represented regions at estimated myBaits® (Arbor BioSciences, MI, USA) hybridization conditions, were removed.

### 2.3 DNA extraction, library preparation and UCE sequence capture

Genomic DNA was extracted from muscle tissue dissected from the pleon or one to several walking legs in the case of large-bodied specimens, from an entire set of legs, or from the whole body in the case of small-bodied specimens (e.g., Peracarida, Branchiopoda). For fresh specimens (collected at most 20 years ago, preserved in *ca*. 95% EtOH, and preserved at -20 °C) we used the Qiagen DNeasy Blood and Tissue kit following the manufacturer’s protocol, with final elution between 100 and 200 µL in ddH_2_O, depending on the amount of starting material. For older museum specimens (collected more than 10 years ago, preserved in 70-80% EtOH and stored at room temperature), we followed the protocol of Tin et al. (2014) for DNA extraction from degraded historical specimens using silica-based magnetic beads, with some in-lab modifications (Derkarabetian et al., 2019), and a final elution volume between 20 and 70 µL in ddH_2_O. These samples are below referred to as “historical” samples. All extractions were quantified on a Qubit 2.0 fluorometer using a dsDNA High Sensitivity kit (Life Technologies, Inc.).

Libraries were prepared using the KAPA HyperPlus kit, following the manufacturer’s protocol with some in-lab modifications (in parts described in Derkarabetian et al., 2019; Moles & Giribet, 2021), particularly for the historical samples. We used up to 250 ng of DNA as input material, however, input from historical samples was usually much lower (down to 4 ng). Fresh samples, as well as more recently collected historical samples were enzymatically fragmented to a target length of 500–700 bp, using 5 µL KAPA Fragmentation Enzyme, 2.5 µL KAPA Fragmentation Buffer (10X) and 17.5 µL DNA sample for a final volume of 25 µL, with incubation times between 3 and 8 min at 37 °C. Fragmentation times for samples of different age and DNA content were optimized by visualizing fragmentation results on an Agilent 2200 TapeStation. Older historical samples did not require fragmentation as they were naturally degraded, and 25 µL of the eluted DNA went directly into end-repair and A-tailing. End-repair and A-tailing was conducted using the KAPA HyperPlus enzyme mix for fresh, enzymatically fragmented samples, and KAPA HyperPrep enzyme mix for historical, naturally fragmented samples, with 30 min incubation at 65 °C. This step was immediately followed by adapter ligation, using 10 µM universal iTru stubs and 45 min incubation at 20 °C for fresh, high-input samples, and 5 µM stubs and up to 60 min incubation at 20 °C for historical and low-input samples. A post-ligation cleanup was carried out using freshly prepared Serapure SpeedBeads (Rohland & Reich, 2012) with 0.8X beads for fresh, and up to 3X beads for old and low-input samples. Fifteen µL of ligated libraries were used in library amplification, with 25 µL 2X KAPA HiFi HotStart ReadyMix, 5 µL individual i5/i7 dual indexing adaptors (Glenn et al., 2019), and the following thermal protocol: 45 s at 98 °C, 10–18 cycles of 15 s at 98 °C, 30 s at 60 °C, 60 s at 72 °C (number of cycles adjusted to Qubit measures of post-ligation DNA concentration), and 5 min at 72 °C final extension. Amplified libraries were purified with SpeedBeads (1X for high-, 3X for low-input samples), quantified, and pooled into equimolar batches of eight samples with 250 ng DNA per sample. If necessary, pools were speed-vacuumed to a final volume of 14 µL.

Hybridization followed the myBaits® Hybridization Capture for Targeted NGS manual v 5.01 with the following modifications: Hybridization time for pools of fresh samples was 24 h at 60 °C, for pools of historical and low-input samples we used a touchdown-protocol with 4 h at 62 °C, 16 h at 60 °C and 4 h at 55 °C. This decreases hybridization specificity, but increases hybridization yield for degraded samples. Fifteen µL of hybridized pools were amplified for 14– 18 cycles using the same thermal protocol as described above, purified with AMPure beads (1.8X for pools of high-, 3X for pools of low-input samples), quantified on a Qubit 2.0, and size estimated on an Agilent TapeStation 2200. A final 1X bead cleanup was performed on pools where adapter-dimers were present. Amplified, hybridized pools were pooled in equimolar amounts and sequenced on an Illumina NovaSeq platform (paired-end, 150 bp) at the Bauer Core Facility, Faculty of Arts and Sciences, Harvard University.

### 2.4 Bioinformatics and phylogenetic analyses

Raw Illumina reads for the Malacostraca dataset were demultiplexed and processed using PHYLUCE version 1.7.2 following the workflow in the online tutorial. Adapters and low-quality bases were removed with illumiprocessor (Faircloth, 2013), which implements trimmomatic (Bolger et al., 2014). Contigs were assembled with SPAdes version 3.15.5 (Prjibelski et al., 2020) using the “--careful” option to reduce the number of mismatches and indels. Probes were matched to the assembled contigs with phyluce_assembly_match_contigs_to_probes, with a 65% threshold value for minimum locus coverage and identity. UCE sequences from all taxa were extracted to individual FASTA files per locus, including incomplete loci that were recovered only in a subset of the taxa. At this step, we also included contigs from the 22 genomes used for probe set design and *in-silico* tests, “harvested” with phyluce_probe_slice_sequence_from_genomes (see Table S1 for UCE recovery summary statistics for these samples). Extracted sequences were aligned with MAFFT (Katoh & Standley, 2013), and alignments were trimmed with GBlocks (Castresana, 2000; Talavera & Castresana, 2007), using very conservative settings, i.e. -b1 0.5, - b2 0.85, -b3 4, -b4 8, suitable for phylogenetic analyses on high taxonomic levels. We built a 50% taxon coverage matrix, and carried out a further cleaning step with CIAlign (Tumescheit et al., 2022), to crop long gaps near sequence ends (threshold --crop_ends_mingap_perc 0.02) and to remove paralogous and outlier sequences (threshold --remove_divergent_minperc 0.65). All alignments with historical samples were additionally inspected manually in Geneious Prime version 2023.0.1 (Kearse et al., 2012), further removing non-orthologous sequences and potential contaminations.

Phylogenies for the Malacostraca dataset were estimated using the concatenated 50% taxon coverage matrix in IQ-TREE version 2.1.0 (Minh et al., 2020), with model selection using ModelFinder (Kalyaanamoorthy et al., 2017) and ultrafast bootstrap (Hoang et al., 2018) with 1500 replicates. We ran IQ-TREE on both unpartitioned and partitioned matrices, in the latter case using the implemented terrace aware approach (Chernomor et al., 2016) and PartitionFinder (Lanfear et al., 2012). The resulting consensus trees were visualized with iTOL version 6.7.4 (Letunic & Bork, 2021), and edited in Inkscape version 1.2.

Assembly, UCE extraction and alignment for the *Eumunida* and *Aratus* datasets followed the same pipeline as above. We used less conservative trimming settings for GBlocks (--b1 0.5 -- b2 0.5 --b3 6 --b4 6 for the *Eumunida* dataset, --b1 0.5 --b2 0.5 --b3 10 --b4 4 for the *Aratus* dataset, respectively), but a slightly higher threshold for outlier removal with CIAlign (-- remove_divergent_minperc 0.75), reflecting the shallower taxonomic level of these datasets. In addition to assembly and UCE recovery statistics for the *Eumunida* and *Aratus* datasets, we assessed recovery of genetic variation on species- and population-level via “smilograms’’ using the PHYLUCE function phyluce_align_get_smilogram_from_alignments on the full *Aratus* dataset, and a subset of the *Eumunida* dataset including eight samples representing five *Eumunida* species from the Western Indian Ocean (*E. bispinata*, *E. minor*, *E. multispina*, *E. similior* and *E. spiridonovi*). “Smilograms” visualize the frequency of base variations across UCE loci in relation to the distance from the alignment center, usually showing increasing variability in the flanking regions compared to the UCE core.

## 3 Results

### 3.1 UCE recovery and probe set efficacy

The final probe set after filtering and final testing included 20,304 probes targeting 1,384 loci.

For the Malacostraca dataset, trimmed Illumina reads of the *in-vitro* samples were assembled into an average of 227,094 contigs per sample (range 1,982–1,108,273). Average raw UCE locus recovery per sample was 632 across all samples (range 5–1,166), with the oldest sample in the dataset, *Aratus pisonii* collected in 1865, recovering the lowest number of loci. When considering only fresh samples (collection date 2012 or later), average raw UCE locus recovery was 785 (range 243–1166). Average locus length across all *in-vitro* Malacostraca samples was 829 bp (range of mean locus length per sample 213–1,627 bp), again with the oldest sample having the shortest average UCE length (see Table 1 for detailed assembly and UCE extraction statistics of the *in-vitro* samples). Across malacostracan orders, locus recovery from fresh and *in-silico* samples was generally highest within Decapoda, however, the probe set still recovered > 800 loci from Euphausiacea and Stomatopoda, 454–1015 loci from various Peracarida, and even 381 and 452 loci from the non-malacostracan Copepoda (*Eurytemora affinis*) and Branchiopoda (*Artemia franciscana*), respectively, which were included as outgroups (Table 1, Table S1).

For the *Eumunida* dataset, SPAdes assembly of the trimmed Illumina reads resulted in an average of 124,439 contigs (range 95–745,615). Average raw UCE locus recovery was 587 across all samples (range 8–1,149). We were able to recover 8 loci from the oldest *Eumunida* sample collected in 1906, however, three more recently collected specimens had ten or less raw UCE loci recovered. Average locus length across all samples was 581 bp (range of mean locus length per sample 190–1,339 bp), again with the oldest sample having the shortest average locus length (Table S2).

Lastly, the 19 samples in the *Aratus pisonii* dataset yielded an average of 813,906 assembled contigs from the trimmed Illumina reads (range 1,674–1,671,386). Average raw UCE locus recovery was 609 (range 8–1,017), of which on average 481 loci were retained in the final 50% taxon coverage matrix (range 2–793). When considering only the fresh samples in this dataset (all collected in 2021), average raw locus recovery was 950 (range 865–1,017), and average mean UCE locus length per sample was 670 bp (range 213–1,622 bp). See Table S3 for detailed assembly and UCE extraction summary statistics for this dataset.

Across all three datasets, UCE locus recovery and average locus length clearly decreased with increasing sample age (Figure 1). Still, the probe set recovered 5–26 loci from the three oldest specimens (*Aratus pisonii*, MCZ IZ 6194 and MCZ IZ 6197, collected 164 and 158 years ago, respectively), of which 3–14 loci were retained in the final matrices after trimming and manual inspection. Average UCE lengths for these oldest samples were likewise among the lowest across all samples, ranging between 213 and 262 bp (compare Tables 1, S2, S3). At the same time, UCE recovery and average length showed considerable variation across the more recent historical, as well as the fresh specimens, reflecting differences in specimen preservation and storage (see Discussion for details).

**Figure 1.**
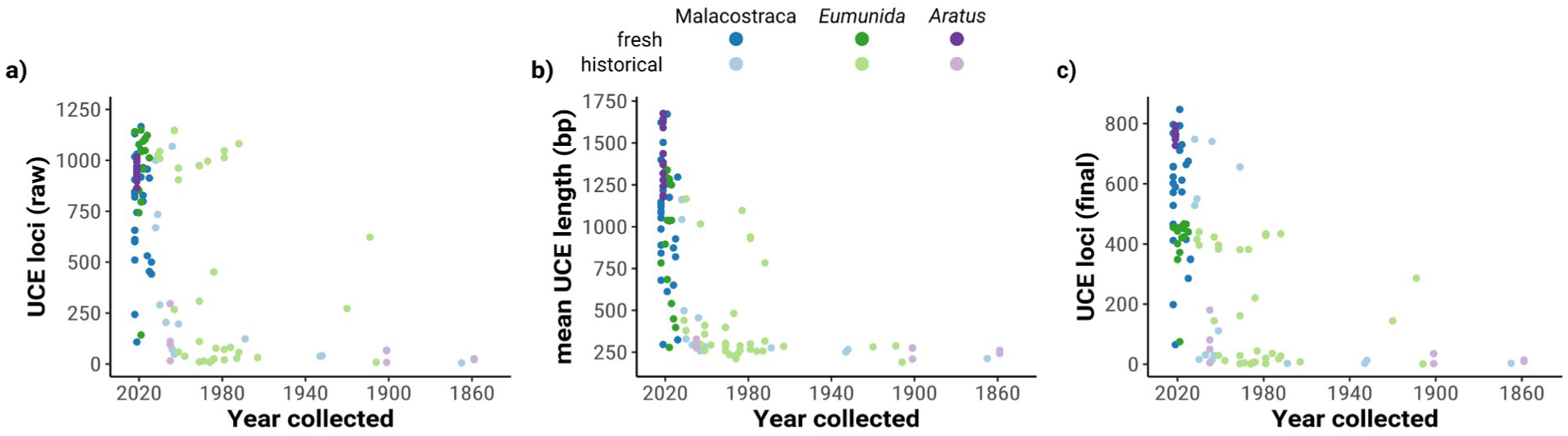
UCE locus recovery and mean UCE locus length for fresh and historical samples across the three datasets used in this study. a) Number of raw (unaligned) UCE loci recovered by PHYLUCE. b) Mean raw UCE length per sample. c) Number of loci in final 50% taxon coverage matrices. Note the different scales on the y-axis

Further exploration of the *Eumunida* and *Aratus* datasets indicated that the targeted UCE loci are also informative at species and population levels. The trimmed and revised 50% taxon coverage matrix of Western Indian Ocean *Eumunida* spp. (five species, eight samples, see Table S2) included 4,016 informative sites at a total length of 316,476 bp across 473 loci. The trimmed and revised 50% taxon coverage matrix of the *Aratus* dataset (four populations, 19 samples, see Table S3) included 18,705 informative sites at a total length of 1,049,132 bp across 813 loci. Sequence variation generally increased with increasing distance from the UCE core region, showing a bimodal/W-shaped distribution for the *Eumunida* subset (Figure 2a) and a more typical, U-shaped distribution for the *Aratus* dataset (Figure 2b).

**Figure 2.**
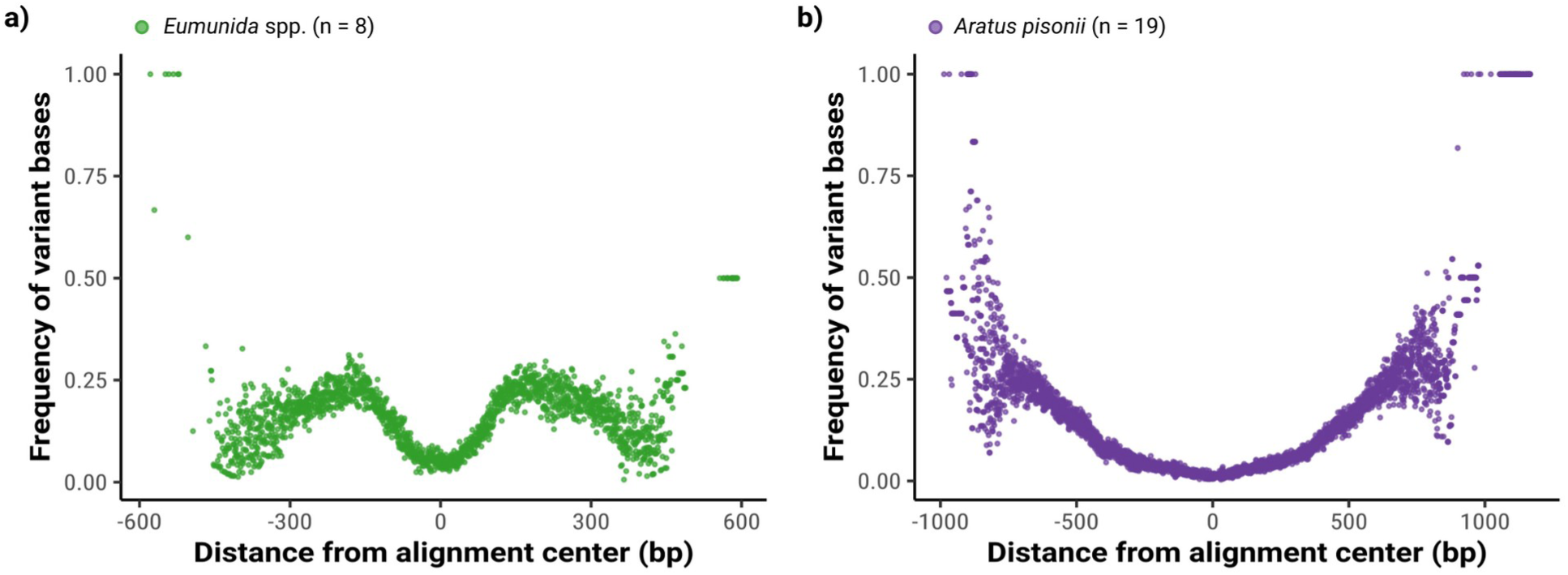
Species- and population-level genetic variation captured by the UCE probe set, shown as the frequency of variant bases in relation to their distance from the center of alignment. Variation data extracted from trimmed 50% taxon coverage matrices. a) “Smilogram” for a subset of five *Eumunida* species from the western Indian Ocean. b) “Smilogram” for a dataset of 23 *Aratus pisonii* specimens collected at four sites around Miami, FL

### 3.2 Phylogenetic analyses

The final, trimmed and revised 50% taxon coverage matrix of the Malacostraca dataset contained 897 UCE loci with an average length of 230 ± 8 bp per locus. The average number of loci per sample in this final matrix was 527 (range 3–863, Table 1 and Table S1). The concatenated alignment had a length of 206,250 bp and contained 100,102 informative sites, with an average of 112 informative sites per locus. Maximum-likelihood analyses of the final dataset recovered a fully resolved phylogeny of the included taxa (Figure 1). Node support was generally high, with exceptions for some of the deeper nodes, and/or in association with clades containing historical samples. As the tree topologies for the unpartitioned and partitioned ML analyses were identical, we present only the tree based on the unpartitioned analysis in the main manuscript, and the tree based on the partitioned analysis as supplementary material (Figure S1).

*Artemia franciscana* (Branchiopoda) was selected as the outgroup of our taxon set. *Eurytemora affinis* (Copepoda) is recovered as the sister group of all the remaining taxa, which represent the Malacostraca. Leptostraca (*Nebalia* sp.) is recovered at the earliest diverging clade, sister group to all other Malacostraca. The superorder Peracarida is not recovered as monophyletic in our analysis. Instead, a clade including all isopods (*Armadillidium*, *Bathynomus* and *Idotea*) groups with Cumacea, while Lophogastrida, represented by *Gnathophausia zoea,* is recovered as an independent lineage, and all amphipods (*Hyaella*, *Parhyale*, *Gammarus* and Maeridae) form a sister clade to the remaining malacostracan orders, i.e., Stomatopoda, Euphausiacea and Decapoda. However, the node splitting the Isopoda/Cumacea clade from the other Peracarida has only low bootstrap support (BS < 70%). Our analysis recovers Decapoda as monophyletic, and Euphausiacea, represented by *Euphausia pacifica*, as their sister group, which is concordant with a superorder Eucarida. Stomatopoda are suggested as the sister group of Eucarida. Within decapods, all infraorders included in our taxon set are recovered as independent lineages. A division between Dendrobranchiata and a monophyletic group containing all other decapod infraorders is strongly supported. This latter group, concordant with Pleocyemata, is divided in two monophyletic clades, one formed by Caridea and Stenopodidae, the other by the ‘reptant’ (crawling/walking) groups, i.e., Achelata, Anomura, Astacidea, Axiidea, Brachyura, and Polychelida. Axiidea clusters with a clade consisting of Astacidea, Achelata and Polychelida, and the two remaining groups, Anomura and Brachyura, form a second monophyletic group of reptant decapods, which is concordant with Meiura. Within the Brachyura, our analysis supports an early diverging position of the Dromioidea with respect to the other groups, represented by *Metadynomede* sp., and also recovers the two major clades within the Eubrachyura, i.e. Heterotremata, represented by *Actaeodes, Callinectes, Chionoecetes, Cymo*, *Portunus,* and *Trapezia*; and Thoracotremata, represented by *Aratus, Armases, Eriocheir, Gecarcinus, Pachygrapsus,* and *Percnon* (Figure 1).

The historical samples included in the dataset were mostly placed at their expected positions in the phylogeny, despite a strong decrease of the number of retained loci with sample age (compare Table 1, Figure 1). The historical *Aratus pisonii, Armadillidium vulgare, Artemia franciscana, Hyalella azteca* and *Penaeus japonicus* samples clustered with their conspecific fresh or *in-silico* samples, respectively. Unequal branch lengths between several of these sample pairs are likely caused by the natural DNA degradation in the historical samples. The historical *Callinectes sapidus* sample was recovered within Portunidae, while *Eriocheir sinensis* clustered with the sesarmid samples (*Aratus, Armases*) rather than the conspecific *in-silico* sample. Notably, the almost 20 year-old *Nebalia* sample, the only representative of Leptostraca in our analysis, was recovered as the sister group to all other Malacostraca, as expected.

## 4 Discussion

In this study, we present a newly designed UCE probeset specifically tailored for Malacostraca, with a particular focus on Decapoda and illustrate its application and effectiveness at different evolutionary scales. Over the past few years, UCEs have emerged as valuable markers for phylogenomic reconstruction and population studies due to their ability to provide large numbers of homologous loci. While UCE probe sets have been developed for many major taxa (Faircloth et al., 2012, e.g. 2013; Goulding et al., 2023; Moles & Giribet, 2021; Quattrini et al., 2018; Smith et al., 2014; Starrett et al., 2017), no probe set was available for the most diverse marine invertebrate class: Malacostraca. Our research addresses this gap by introducing a specialized UCE probeset for investigating the intricate diversity within Malacostraca, with a specific emphasis on the diverse order Decapoda.

We identified almost 1,400 highly conserved genomic regions between taxa of the hyperdiverse Malacostraca, and designed about 20,000 probes to target these UCEs. We tested the efficacy of this new genomic resource on three datasets representing various levels of phylogenetic depth. The mean number of UCE loci recovered from our main Malacostraca dataset of 31 fresh, 10 historical and 21 genome-derived *in-silico* samples (743 raw and 621 final loci in the 50% matrix, respectively), as well as mean locus length (742 bp in the raw, and 230 bp in the 50% matrix after GBlocks trimming, respectively), and the proportion of informative sites in the final alignment (48.5%) are comparable to similar-sized UCE probe sets for other invertebrate groups (compare e.g., Moles and Giribet, 2021; Quattrini et al., 2018; Starrett et al., 2017). It should be noted that we achieved these recovery statistics with a comparatively high proportion of historical samples in the dataset. When including fresh and *in-silico* samples only, locus recovery increases to a mean of 853 loci per sample in the raw, and 621 loci per sample in the final 50% taxon coverage matrix.

All three datasets analyzed in this study demonstrate the capability of the new probe set to capture sequence data from historical specimens, regularly highlighted as a key asset of UCEs in studies targeting various invertebrate and vertebrate taxa (e.g. Blaimer et al., 2016; Derkarabetian et al., 2019; Moles & Giribet, 2021; Salter et al., 2022). We were able to recover up to 25 loci from specimens collected in 1859 (i.e., 164 years old), which are to our knowledge among the oldest used in targeted UCE sequencing studies (compare Derkarabetian et al., 2019), up to 68 loci from specimens collected in 1901, and as much as 242 loci from a specimen collected in 1920 (see Table S2, S3). While the number of loci from historical specimens dropped considerably during alignment trimming and revisions, the retained loci were still informative enough to put most historical specimens to their expected phylogenetic position (Figure 3). At the same time, we observed very low locus recovery in a few samples collected less than 20 years ago (e.g. *Callinectes sapidus*, MCZ IZ 23424, coll. 2010: 14 loci in final matrix, or *Aratus pisonii* UF 11375, coll. 2005: 5 loci in final matrix). A likely cause for the apparent lack of a clearer correlation between sample age and locus recovery are vastly varying collection, preservation, and storage histories among collection events of “standard” museum specimens, which are still mostly focused on preserving external morphology. Particularly in hard-bodied crustaceans like crabs, soft tissues usually used for DNA extraction may quickly degrade due to slow ethanol penetration through the exoskeleton, if it is not manually perforated (own observations). As fixation and storage histories of specimens are oftentimes unknown and/or not reported in collection databases, the use of historical specimens for sequencing experiments will inevitably include the risk of selecting specimens that are unsuitable for molecular work (compare e.g., Bernstein & Ruane, 2022; Wandeler et al., 2007).

**Figure 3.**
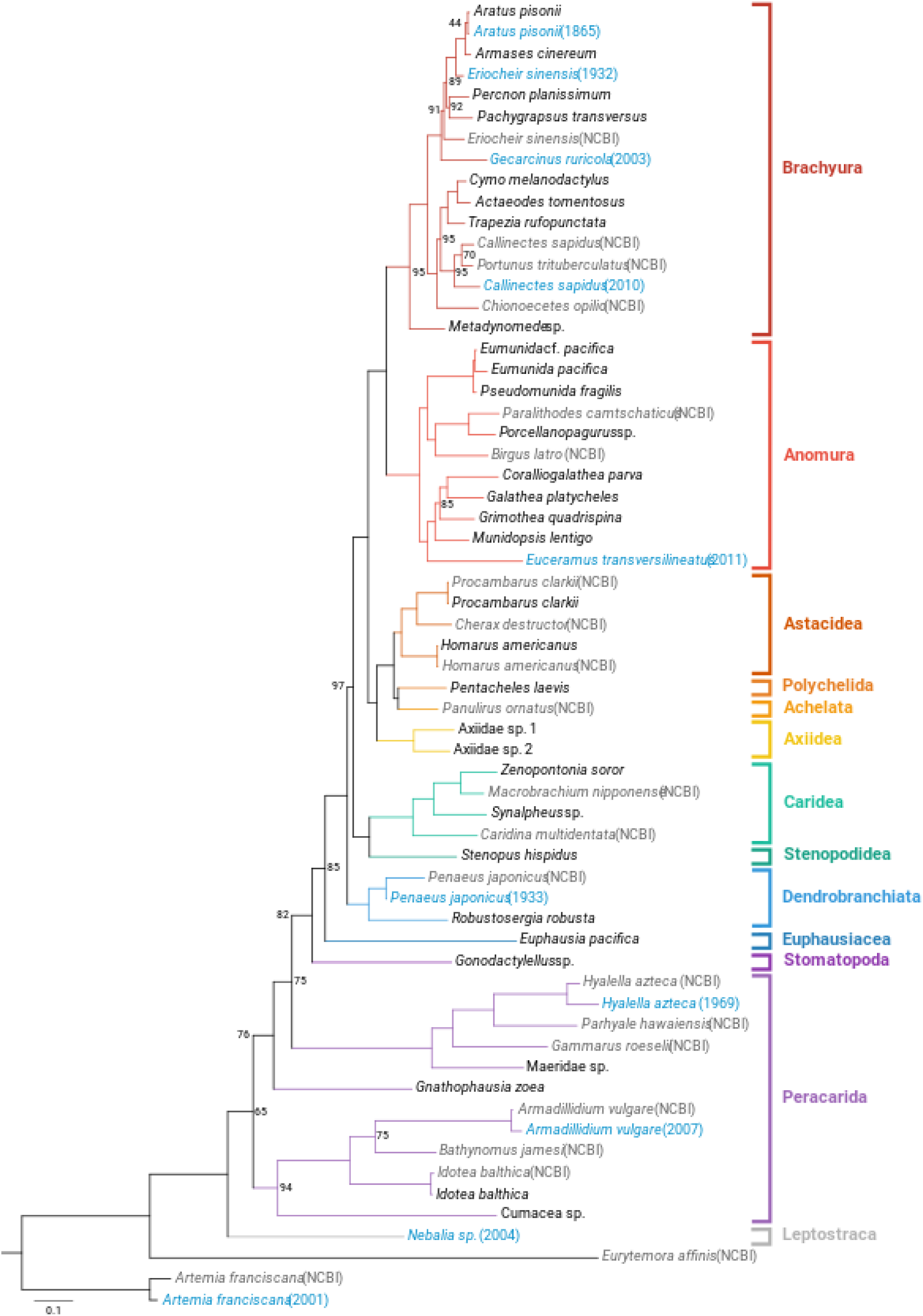
Phylogenetic hypothesis (maximum likelihood) for Malacostraca, based on a 50% taxon-coverage matrix of 897 UCE loci. Historical samples are indicated in blue, with collection years in parenthesis. *In-silico* samples (NCBI) are indicated in gray. Species used for probe set design are in bold. Bootstrap support values are given for nodes with support < 100%

Our data furthermore show a notable universal applicability of the probe set across, and even beyond, Malacostraca. With the probe set design strongly focused on Decapoda, we consequently recovered the highest numbers of loci from samples within this group (919–1,196 loci, i.e. 66.4–86.4% of the total probe set for the *in-silico* samples). However, between 450 and > 1,000 loci were recovered from fresh or *in-silico* Euphausiacea, Stomatopoda, and several Peracarida, and still between about 200 and 450 loci from fresh and *in-silico* Copepoda and Branchiopoda. Thus, the probe set performs well across a vastly divergent group, spanning at least 540 million years of evolution (compare Bernot et al., 2022). In this respect, the Decapoda probe set appears equivalent to the Arachnida probe set, spanning a similarly divergent lineage (Starrett et al., 2017), corroborating Bossert and Danforth’s (2018) findings on the universal character of UCEs. At the other end of the spectrum of phylogenetic depth, the genus-(*Eumunida* spp.) and species-level (*Aratus pisonii*) datasets show increasing sequence variability in UCE locus alignments with increasing distance from the UCE core region, as expected (compare Figure 2). While the proportion of variable sites in the final alignments of these datasets is relatively low (1.3% and 1.8% for the *Eumunida* and *Aratus* datasets, respectively), our data still clearly indicate the utility of the probe set for e.g., biogeographic or population genomic studies on genus- and species-level datasets. Optimizing locus trimming and filtering settings towards keeping longer flanking regions in the final alignments could be a way to improve the recovery of variable sites. Another way to increase overall locus recovery and contig length could be to combine contigs derived from different assemblers (e.g. ABySS and SPAdes) before extracting UCE loci, as has been demonstrated in UCE studies on Opiliones and heterobranch molluscs (Derkarabetian et al., 2019; Moles & Giribet, 2021).

The UCE-based malacostracan phylogeny demonstrates the utility of our probe set across several phylogenetic levels. Overall, the tree topology agrees well with recent malacostracan phylogenies based on transcriptomes (Bernot et al., 2022) and mitogenomes (Höpel et al., 2022). Specifically, the focal group for probe set design, Decapoda, is recovered as monophyletic with full support, and most of the internal relationships within the order are congruent with Wolfe et al.’s (2019) phylogeny based on AHE data. The only major discrepancy to previously published decapod phylogenies is the clustering of Axiidea in a clade with Astacidea, Polychelida and Achelata, instead of clustering with Meiura (Anomura + Brachyura) (e.g. Shen et al., 2015; Wolfe et al., 2019). The historical samples included in the Malacostraca dataset are all recovered at their expected positions in the phylogeny, with the exception of the almost 100 years-old sample of *Eriocheir sinensis*, which did not cluster with its conspecific *in-silico* sample, but the closely related sesarmid crabs, an artifact likely caused by low locus recovery and high DNA degradation of this historical sample. Regarding the deeper malacostracan phylogeny, the UCE loci recover Leptostraca as the sister group to the remaining Malacostraca, as well as a clade including Stomatopoda, Euphausiacea and Decapoda, in line with the findings of Bernot et al. (2022) and Höpel et al. (2022). Peracarida, on the other hand, are not recovered as monophyletic. While their monophyly has been occasionally disputed (Poore, 2005; Richter & Scholtz, 2001, and references therein), morphological and multilocus phylogenies, as well as mitogenomic and phylotranscriptomic studies support a monophyletic Peracarida (Bernot et al., 2022; Höpel et al., 2022; Schwentner et al., 2018; Spears et al., 2005). A potential explanation for lacking monophyly of Peracarida in our data could be insufficient taxon sampling within the group (only one sample from each Cumacea and Lophogastrida, and no representatives of Mysida, Tanaidacea, and Thermosbaenacea), and strong taxon sampling bias towards Decapoda (particularly Anomura and Brachyura) in the Malacostraca dataset (compare Bernot et al., 2022). Furthermore, the fact that Peracarida were represented only by one amphipod genome in our probe set design might have caused a lack of loci that can resolve the deeper nodes within this group, which is almost as speciose as the Decapoda. We would like to stress, however, that the intention of this study was not to fully resolve phylogenetic relationships among and within malacostracan orders, but rather to provide a proof of concept, and a demonstration of the performance and utility of this new UCE probe set. Future analyses will focus on peracarid relationships to test this clade.

Overall, our data provide strong evidence for the versatility of the UCE probe set we present in this study, both within the focal group of probe set design for Decapoda, as well as across the highly divergent Malacostraca. They highlight the universality of the targeted probes and their ability to recover deep phylogenetic relationships, and genetic variation from taxonomically shallow datasets alike. They furthermore demonstrate the utility of the probe set to extract informative sequence data from historical museum specimens. This is particularly beneficial for studies including rare, or difficult to sample species, such as deep-sea crustaceans, as many of them will only be available as historical museum samples. Therefore, we expect our probe set to become a valuable and affordable resource for targeted sequencing studies in Malacostraca across multiple taxonomic levels and along varying lines of research, providing an alternative complement to the AHE probes published by Wolfe et al. (2019). This probeset should therefore allow researchers to explore a wide range of evolutionary and population studies within one of the most diverse and economically significant invertebrate taxa.

## Acknowledgements

Funding for this study was provided by a Walter-Benjamin postdoctoral fellowship from the Deutsche Forschungsgemeinschaft (DFG, German Research Foundation) to J.C.G. (project number 447933028), by a Biodiversity Postdoctoral Fellowship from the Museum of Comparative Zoology, Harvard University (MCZ) to P.C.R.F., as well as by Putnam Expedition Grants from the MCZ to J.C.G. and P.C.R.F. for collecting specimens in Florida and on Guam. Lab work and sequencing were financially supported by internal funds from the MCZ. We thank Claire Hartmann (Reardon) of the Bauer Core Facility at Harvard University for her great help in troubleshooting and optimizing sequencing results. Hector Torrado and David Combosch supported us during field work on Guam, as did Linnea and Sinikka Lennartz during field work in Florida. The historical specimens used in this study were provided by the Museum of Comparative Zoology, the Musée national de l’Histoire naturelle, the Smithsonian National Museum of Natural History, the Florida Museum of Natural History, and the California Academy of Sciences, and we thank all curators who generously granted our loan and sampling requests including Laure Corbari, Paula Martin-Lefevre, Martha Nizinski, Gustav Paulay, Christina Piotrowski, and, Karen Reed. P.C.R.F. is grateful for Laure Corbari’s and Paula Martin-Lefevre’s hospitality during her stays at the crustacean collection of the MNHN in Paris.

## Data Accessibility and Benefit-Sharing

Raw Oxford Nanopore reads of *Aratus pisonii* used for probe set design, as well as demultiplexed Illumina raw reads from the hybridization pools have been deposited in the SRA (BioProject PRJNA988117). The probe set, the final UCE locus alignment matrices, and the resulting phylogenies are available from the Harvard Dataverse (https://doi.org/10.7910/DVN/WASGMF).

## Author contributions

J.C.G. designed the research with support from G.G. and S.D., J.C.G. and P.C.R.F. performed the research with support from S.D., G.G. provided financial support for reagents and sequencing, J.C.G. and P.C.R.F. wrote the paper, and all authors edited and approved the final manuscript.

## Supporting information

The following supporting information is provided in a separate document:

**Table S1** Accession information and UCE locus recovery for the *in-silico* samples included in the Malacostraca dataset

**Table S2** Assembly and UCE extraction summary statistics for the *Eumunida* dataset

**Table S3** Assembly and UCE summary statistics for the *Aratus* dataset

**Figure S1** Maximum-likelihood tree for Malacostraca, based on a partitioned analysis of a 50% taxon coverage matrix

## References

Ballesteros, J. A., Setton, E. V. W., Santibáñez-López, C. E., Arango, C. P., Brenneis, G., Brix, S., Corbett, K. F., Cano-Sánchez, E., Dandouch, M., Dilly, G. F., Eleaume, M. P., Gainett, G., Gallut, C., McAtee, S., McIntyre, L., Moran, A. L., Moran, R., López-González, P. J., Scholtz, G., … Sharma, P. P. (2021). Phylogenomic resolution of sea spider diversification through integration of multiple data classes. Molecular Biology and Evolution, 38(2), 686–701. https://doi.org/10.1093/molbev/msaa228

Bejerano, G., Pheasant, M., Makunin, I., Stephen, S., Kent, W. J., Mattick, J. S., & Haussler, D. (2004). Ultraconserved elements in the human genome. Science, 304(5675), 1321–1325. https://doi.org/10.1126/science.1098119

Bernot, J. P., Owen, C. L., Wolfe, J. M., Meland, K., Olesen, J., & Crandall, K. A. (2022). *Major revisions in pancrustacean phylogeny with recommendations for resolving challenging nodes* [Preprint]. Evolutionary Biology. https://doi.org/10.1101/2022.11.17.514186

Bernstein, J. M., & Ruane, S. (2022). Maximizing molecular data from low-quality fluid-preserved specimens in natural history collections. Frontiers in Ecology and Evolution, 10. https://www.frontiersin.org/articles/10.3389/fevo.2022.893088

Blaimer, B. B., Lloyd, M. W., Guillory, W. X., & Brady, S. G. (2016). Sequence capture and phylogenetic utility of genomic ultraconserved elements obtained from pinned insect specimens. PLOS ONE, 11(8), e0161531. https://doi.org/10.1371/journal.pone.0161531

Bolger, A. M., Lohse, M., & Usadel, B. (2014). Trimmomatic: A flexible trimmer for Illumina sequence data. Bioinformatics, 30(15), 2114–2120. https://doi.org/10.1093/bioinformatics/btu170

Bossert, S., & Danforth, B. N. (2018). On the universality of target-enrichment baits for phylogenomic research. Methods in Ecology and Evolution, 9(6), 1453–1460. https://doi.org/10.1111/2041-210X.12988

Boyer, S. L., Dohr, S. R., Tuffield, M. S., Shu, Y., Moore, C. D., Hahn, K. M., Ward, R. S., Nguyen, P., Morisawa, R., Boyer, S. L., Dohr, S. R., Tuffield, M. S., Shu, Y., Moore, C. D., Hahn, K. M., Ward, R. S., Nguyen, P., & Morisawa, R. (2022). Diversity and distribution of the New Zealand endemic mite harvestman genus *Aoraki* (Arachnida, Opiliones, Cyphophthalmi, Pettalidae), with the description of two new species. Invertebrate Systematics, 36(4), 372–387. https://doi.org/10.1071/IS21044

Castresana, J. (2000). Selection of conserved blocks from multiple alignments for their use in phylogenetic analysis. Molecular Biology and Evolution, 17(4), 540–552. https://doi.org/10.1093/oxfordjournals.molbev.a026334

Chernomor, O., von Haeseler, A., & Minh, B. Q. (2016). Terrace aware data structure for phylogenomic inference from supermatrices. Systematic Biology, 65(6), 997–1008. https://doi.org/10.1093/sysbio/syw037

Cowman, P. F., Quattrini, A. M., Bridge, T. C. L., Watkins-Colwell, G. J., Fadli, N., Grinblat, M., Roberts, T. E., McFadden, C. S., Miller, D. J., & Baird, A. H. (2020). An enhanced target-enrichment bait set for Hexacorallia provides phylogenomic resolution of the staghorn corals (Acroporidae) and close relatives. Molecular Phylogenetics and Evolution, 153, 106944. https://doi.org/10.1016/j.ympev.2020.106944

de Miranda, G. S., Kulkarni, S. S., Tagliatela, J., Baker, C. M., Giupponi, A. P. L., Labarque, F. M., Gavish-Regev, E., Rix, M. G., Carvalho, L. S., Fusari, L. M., Wood, H. M., & Sharma, P. P. (2022). The rediscovery of a relict unlocks the first global phylogeny of whip spiders (Amblypygi) (p. 2022.04.26.489547). bioRxiv. https://doi.org/10.1101/2022.04.26.489547

Derkarabetian, S., Benavides, L. R., & Giribet, G. (2019). Sequence capture phylogenomics of historical ethanol-preserved museum specimens: Unlocking the rest of the vault. Molecular Ecology Resources, 19(6), 1531–1544. https://doi.org/10.1111/1755-0998.13072

Derkarabetian, S., Paquin, P., Reddell, J., & Hedin, M. (2022). Conservation genomics of federally endangered *Texella* harvester species (Arachnida, Opiliones, Phalangodidae) from cave and karst habitats of central Texas. Conservation Genetics, 23(2), 401–416. https://doi.org/10.1007/s10592-022-01427-9

Derkarabetian, S., Starrett, J., Tsurusaki, N., Ubick, D., Castillo, S., & Hedin, M. (2018). A stable phylogenomic classification of Travunioidea (Arachnida, Opiliones, Laniatores) based on sequence capture of ultraconserved elements. ZooKeys, 760, 1–36. https://doi.org/10.3897/zookeys.760.24937

Faircloth, B. C. (2013). Illumiprocessor: A trimmomatic wrapper for parallel adapter and quality trimming. J9ILL.

Faircloth, B. C. (2016). PHYLUCE is a software package for the analysis of conserved genomic loci. Bioinformatics, 32(5), 786–788. https://doi.org/10.1093/bioinformatics/btv646

Faircloth, B. C. (2017). Identifying conserved genomic elements and designing universal bait sets to enrich them. Methods in Ecology and Evolution, 8(9), 1103–1112. https://doi.org/10.1111/2041-210X.12754

Faircloth, B. C., McCormack, J. E., Crawford, N. G., Harvey, M. G., Brumfield, R. T., & Glenn, T. C. (2012). Ultraconserved elements anchor thousands of genetic markers spanning multiple evolutionary timescales. Systematic Biology, 61(5), 717–726. https://doi.org/10.1093/sysbio/sys004

Faircloth, B. C., Sorenson, L., Santini, F., & Alfaro, M. E. (2013). A phylogenomic perspective on the radiation of ray-finned fishes based upon targeted sequencing of ultraconserved elements (UCEs). PLOS ONE, 8(6), e65923. https://doi.org/10.1371/journal.pone.0065923

Glenn, T. C., Nilsen, R. A., Kieran, T. J., Sanders, J. G., Bayona-Vásquez, N. J., Finger, J. W., Pierson, T. W., Bentley, K. E., Hoffberg, S. L., Louha, S., Leon, F. J. G.-D., del Rio Portilla, M. A., Reed, K. D., Anderson, J. L., Meece, J. K., Aggrey, S. E., Rekaya, R., Alabady, M., Belanger, M., … Faircloth, B. C. (2019). Adapterama I: Universal stubs and primers for 384 unique dual-indexed or 147,456 combinatorially-indexed Illumina libraries (iTru & iNext). PeerJ, 7, e7755. https://doi.org/10.7717/peerj.7755

Glon, H., Quattrini, A., Rodríguez, E., Titus, B. M., & Daly, M. (2021). Comparison of sequence-capture and ddRAD approaches in resolving species and populations in hexacorallian anthozoans. Molecular Phylogenetics and Evolution, 163, 107233. https://doi.org/10.1016/j.ympev.2021.107233

Gonzalez, B., Martínez, A., Institute for Water Research, Italian National Research Council, Olesen, J., Truskey, S. B., Ballou, L., Allentoft-Larsen, M., Daniels, J., Heinerth, P., Parrish, M., Manco, N., Ward, J., Iliffe, T. M., Osborn, K. J., & Worsaae, K. (2020). Anchialine biodiversity in the Turks and Caicos Islands: New discoveries and current faunal composition. International Journal of Speleology, 49(2), 71–86. https://doi.org/10.5038/1827-806X.49.2.2316

Goulding, T. C., Strong, E. E., & Quattrini, A. M. (2023). Target-capture probes for phylogenomics of the Caenogastropoda. Molecular Ecology Resources, n/a(n/a). https://doi.org/10.1111/1755-0998.13793

Grüning, B., Dale, R., Sjödin, A., Chapman, B. A., Rowe, J., Tomkins-Tinch, C. H., Valieris, R., & Köster, J. (2018). Bioconda: Sustainable and comprehensive software distribution for the life sciences. Nature Methods, 15(7), Article 7. https://doi.org/10.1038/s41592-018-0046-7

Hedin, M., Derkarabetian, S., Alfaro, A., Ramírez, M. J., & Bond, J. E. (2019). Phylogenomic analysis and revised classification of atypoid mygalomorph spiders (Araneae, Mygalomorphae), with notes on arachnid ultraconserved element loci. PeerJ, 7, e6864. https://doi.org/10.7717/peerj.6864

Hedin, M., Foldi, S., & Rajah-Boyer, B. (2020). Evolutionary divergences mirror Pleistocene paleodrainages in a rapidly-evolving complex of oasis-dwelling jumping spiders (Salticidae, *Habronattus tarsalis*). Molecular Phylogenetics and Evolution, 144, 106696. https://doi.org/10.1016/j.ympev.2019.106696

Hoang, D. T., Chernomor, O., von Haeseler, A., Minh, B. Q., & Vinh, L. S. (2018). Ufboot2: Improving the ultrafast bootstrap approximation. Molecular Biology and Evolution, 35(2), 518–522. https://doi.org/10.1093/molbev/msx281

Höpel, C. G., Yeo, D., Grams, M., Meier, R., & Richter, S. (2022). Mitogenomics supports the monophyly of Mysidacea and Peracarida (Malacostraca). Zoologica Scripta, 51(5), 603– 613. https://doi.org/10.1111/zsc.12554

Hou, Z., Sket, B., & Li, S. (2014). Phylogenetic analyses of Gammaridae crustacean reveal different diversification patterns among sister lineages in the Tethyan region. Cladistics, 30(4), 352–365. https://doi.org/10.1111/cla.12055

Huang, W., Li, L., Myers, J. R., & Marth, G. T. (2012). ART: A next-generation sequencing read simulator. Bioinformatics, 28(4), 593–594. https://doi.org/10.1093/bioinformatics/btr708

Kalyaanamoorthy, S., Minh, B. Q., Wong, T. K. F., von Haeseler, A., & Jermiin, L. S. (2017). ModelFinder: Fast model selection for accurate phylogenetic estimates. Nature Methods, 14(6), Article 6. https://doi.org/10.1038/nmeth.4285

Katoh, K., & Standley, D. M. (2013). Mafft multiple sequence alignment software version 7: Improvements in performance and usability. Molecular Biology and Evolution, 30(4), 772–780. https://doi.org/10.1093/molbev/mst010

Kearse, M., Moir, R., Wilson, A., Stones-Havas, S., Cheung, M., Sturrock, S., Buxton, S., Cooper, A., Markowitz, S., Duran, C., Thierer, T., Ashton, B., Meintjes, P., & Drummond, A. (2012). Geneious Basic: An integrated and extendable desktop software platform for the organization and analysis of sequence data. Bioinformatics, 28(12), 1647–1649. https://doi.org/10.1093/bioinformatics/bts199

Kent, W. J. (2002). BLAT—The BLAST-Like Alignment Tool. Genome Research, 12(4), 656– 664. https://doi.org/10.1101/gr.229202

Lanfear, R., Calcott, B., Ho, S. Y. W., & Guindon, S. (2012). Partitionfinder: Combined selection of partitioning schemes and substitution models for phylogenetic analyses. Molecular Biology and Evolution, 29(6), 1695–1701. https://doi.org/10.1093/molbev/mss020

Lemmon, A. R., Emme, S. A., & Lemmon, E. M. (2012). Anchored hybrid enrichment for massively high-throughput phylogenomics. Systematic Biology, 61(5), 727–744. https://doi.org/10.1093/sysbio/sys049

Letunic, I., & Bork, P. (2021). Interactive Tree Of Life (iTOL) v5: An online tool for phylogenetic tree display and annotation. Nucleic Acids Research, 49(W1), W293–W296. https://doi.org/10.1093/nar/gkab301

Li, H., Handsaker, B., Wysoker, A., Fennell, T., Ruan, J., Homer, N., Marth, G., Abecasis, G., Durbin, R., & 1000 Genome Project Data Processing Subgroup. (2009). The Sequence Alignment/Map format and SAMtools. Bioinformatics (Oxford, England), 25(16), 2078–2079. https://doi.org/10.1093/bioinformatics/btp352

Lunter, G., & Goodson, M. (2011). Stampy: A statistical algorithm for sensitive and fast mapping of Illumina sequence reads. Genome Research, 21(6), 936–939. https://doi.org/10.1101/gr.111120.110

McCormack, J. E., Faircloth, B. C., Crawford, N. G., Gowaty, P. A., Brumfield, R. T., & Glenn, T. C. (2012). Ultraconserved elements are novel phylogenomic markers that resolve placental mammal phylogeny when combined with species-tree analysis. Genome Research, 22(4), 746–754. https://doi.org/10.1101/gr.125864.111

McCormack, J. E., Tsai, W. L. E., & Faircloth, B. C. (2016). Sequence capture of ultraconserved elements from bird museum specimens. Molecular Ecology Resources, 16(5), 1189–1203. https://doi.org/10.1111/1755-0998.12466

Minh, B. Q., Schmidt, H. A., Chernomor, O., Schrempf, D., Woodhams, M. D., von Haeseler, A., & Lanfear, R. (2020). IQ-TREE 2: New models and efficient methods for phylogenetic inference in the genomic era. Molecular Biology and Evolution, 37(5), 1530–1534. https://doi.org/10.1093/molbev/msaa015

Moles, J., & Giribet, G. (2021). A polyvalent and universal tool for genomic studies in gastropod molluscs (Heterobranchia). Molecular Phylogenetics and Evolution, 155, 106996. https://doi.org/10.1016/J.YMPEV.2020.106996

Patel, T., Robert, H., D’Udekem D’Acoz, C., Martens, K., De Mesel, I., Degraer, S., & Schön, I. (2020). Biogeography and community structure of abyssal scavenging Amphipoda (Crustacea) in the Pacific Ocean. Biogeosciences, 17(10), 2731–2744. https://doi.org/10.5194/bg-17-2731-2020

Poore, G. C. B. (2005). Peracarida: Monophyly, relationships and evolutionary success. Nauplius, 13(1), 1–27.

Prjibelski, A., Antipov, D., Meleshko, D., Lapidus, A., & Korobeynikov, A. (2020). Using SPAdes de novo assembler. Current Protocols in Bioinformatics, 70(1), e102. https://doi.org/10.1002/cpbi.102

Quattrini, A. M., Faircloth, B. C., Dueñas, L. F., Bridge, T. C. L., Brugler, M. R., Calixto-Botía, I. F., DeLeo, D. M., Forêt, S., Herrera, S., Lee, S. M. Y., Miller, D. J., Prada, C., Rádis-Baptista, G., Ramírez-Portilla, C., Sánchez, J. A., Rodríguez, E., & McFadden, C. S. (2018). Universal target-enrichment baits for anthozoan (Cnidaria) phylogenomics: New approaches to long-standing problems. Molecular Ecology Resources, 18(2), 281–295. https://doi.org/10.1111/1755-0998.12736

Quinlan, A. R., & Hall, I. M. (2010). BEDTools: A flexible suite of utilities for comparing genomic features. Bioinformatics, 26(6), 841–842. https://doi.org/10.1093/bioinformatics/btq033

Raxworthy, C. J., & Smith, B. T. (2021). Mining museums for historical DNA: Advances and challenges in museomics. Trends in Ecology & Evolution, 36(11), 1049–1060. https://doi.org/10.1016/j.tree.2021.07.009

Richter & Scholtz. (2001). Phylogenetic analysis of the Malacostraca (Crustacea). Journal of Zoological Systematics and Evolutionary Research, 39(3), 113–136. https://doi.org/10.1046/j.1439-0469.2001.00164.x

Rohland, N., & Reich, D. (2012). Cost-effective, high-throughput DNA sequencing libraries for multiplexed target capture. Genome Research, 22(5), 939–946. https://doi.org/10.1101/gr.128124.111

Salter, J. F., Hosner, P. A., Tsai, W. L. E., McCormack, J. E., Braun, E. L., Kimball, R. T., Brumfield, R. T., & Faircloth, B. C. (2022). Historical specimens and the limits of subspecies phylogenomics in the New World quails (Odontophoridae). Molecular Phylogenetics and Evolution, 175, 107559. https://doi.org/10.1016/j.ympev.2022.107559

Schram, F. R., & Abele, L. (1982). The fossil record and evolution of Crustacea. The Biology of Crustacea, 1, 93–147.

Schwentner, M., Combosch, D. J., Pakes Nelson, J., & Giribet, G. (2017). A phylogenomic solution to the origin of insects by resolving crustacean-hexapod relationships. Current Biology, 27(12), 1818–1824.e5. https://doi.org/10.1016/j.cub.2017.05.040

Schwentner, M., Richter, S., Rogers, D. C., & Giribet, G. (2018). Tetraconatan phylogeny with special focus on Malacostraca and Branchiopoda: Highlighting the strength of taxon-specific matrices in phylogenomics. Proceedings of the Royal Society B: Biological Sciences, 285(1885), 20181524. https://doi.org/10.1098/rspb.2018.1524

Shen, X., Tian, M., Yan, B., & Chu, K. (2015). Phylomitogenomics of Malacostraca (Arthropoda: Crustacea). Acta Oceanologica Sinica, 34(2), 84–92. https://doi.org/10.1007/s13131-015-0583-1

Smith, B. T., Harvey, M. G., Faircloth, B. C., Glenn, T. C., & Brumfield, R. T. (2014). Target capture and massively parallel sequencing of ultraconserved elements for comparative studies at shallow evolutionary time scales. Systematic Biology, 63(1), 83–95. https://doi.org/10.1093/SYSBIO/SYT061

Spears, T., DeBry, R. W., Abele, L. G., & Chodyla, K. (2005). Peracarid monophyly and interordinal phylogeny inferred from nuclear small-subunit ribosomal DNA sequences (Crustacea: Malacostraca: Peracarida). Proceedings of the Biological Society of Washington, 118(1), 117–157. https://doi.org/10.2988/0006-324X(2005)118[117:PMAIPI]2.0.CO;2

Starrett, J., Derkarabetian, S., Hedin, M., Bryson, R. W., McCormack, J. E., & Faircloth, B. C. (2017). High phylogenetic utility of an ultraconserved element probe set designed for Arachnida. Molecular Ecology Resources, 17(4), 812–823. https://doi.org/10.1111/1755-0998.12621

Streicher, J. W., & Wiens, J. J. (2017). Phylogenomic analyses of more than 4000 nuclear loci resolve the origin of snakes among lizard families. Biology Letters, 13(9), 20170393. https://doi.org/10.1098/rsbl.2017.0393

Swan, J. A., Jamieson, A. J., Linley, T. D., & Yancey, P. H. (2021). Worldwide distribution and depth limits of decapod crustaceans (Penaeoidea, Oplophoroidea) across the abyssal-hadal transition zone of eleven subduction trenches and five additional deep-sea features. Journal of Crustacean Biology, 41(1), 1–13. https://doi.org/10.1093/jcbiol/ruaa102

Talavera, G., & Castresana, J. (2007). Improvement of phylogenies after removing divergent and ambiguously aligned blocks from protein sequence alignments. Systematic Biology, 56(4), 564–577. https://doi.org/10.1080/10635150701472164

Tin, M. M. Y., Economo, E. P., & Mikheyev, A. S. (2014). Sequencing degraded DNA from non-destructively sampled museum specimens for RAD-tagging and low-coverage shotgun phylogenetics. PLoS ONE, 9(5). https://doi.org/10.1371/journal.pone.0096793

Tsang, C. T. T., Schubart, C. D., Chu, K. H., Ng, P. K. L., & Tsang, L. M. (2022). Molecular phylogeny of Thoracotremata crabs (Decapoda, Brachyura): Toward adopting monophyletic superfamilies, invasion history into terrestrial habitats and multiple origins of symbiosis. Molecular Phylogenetics and Evolution, 177, 107596. https://doi.org/10.1016/j.ympev.2022.107596

Tumescheit, C., Firth, A. E., & Brown, K. (2022). CIAlign: A highly customisable command line tool to clean, interpret and visualise multiple sequence alignments. PeerJ, 10, e12983. https://doi.org/10.7717/peerj.12983

Vinciguerra, N. T., Tsai, W. L. E., Faircloth, B. C., & McCormack, J. E. (2019). Comparison of ultraconserved elements (UCEs) to microsatellite markers for the study of avian hybrid zones: A test in *Aphelocoma* jays. BMC Research Notes, 12(1), 456. https://doi.org/10.1186/s13104-019-4481-z

Wandeler, P., Hoeck, P. E. A., & Keller, L. F. (2007). Back to the future: Museum specimens in population genetics. Trends in Ecology and Evolution, 22(12), 634–642. https://doi.org/10.1016/j.tree.2007.08.017

Wolfe, J. M., Ballou, L., Luque, J., Watson-Zink, V. M., Ahyong, S. T., Barido-Sottani, J., Chan, T.-Y., Chu, K. H., Crandall, K. A., Daniels, S. R., Felder, D. L., Mancke, H., Martin, J. W., Ng, P. K. L., Ortega-Hernández, J., Theil, E. P., Pentcheff, N. D., Robles, R., Thoma, B. P., … Bracken-Grissom, H. D. (2023). Convergent adaptation of true crabs (Decapoda: Brachyura) to a gradient of terrestrial environments (p. 2022.12.09.519815). bioRxiv. https://doi.org/10.1101/2022.12.09.519815

Wolfe, J. M., Breinholt, J. W., Crandall, K. A., Lemmon, A. R., Lemmon, E. M., Timm, L. E., Siddall, M. E., & Bracken-Grissom, H. D. (2019). A phylogenomic framework, evolutionary timeline and genomic resources for comparative studies of decapod crustaceans. Proceedings of the Royal Society B: Biological Sciences, 286(1901). https://doi.org/10.1098/rspb.2019.0079

Zhang, Y. M., Williams, J. L., & Lucky, A. (2019). Understanding UCEs: A comprehensive primer on using ultraconserved elements for arthropod phylogenomics. Insect Systematics and Diversity, 3(5). https://doi.org/10.1093/ISD/IXZ016

