## Supporting information for "From the shallows to the depths: A new probe set to target ultraconserved elements for Malacostraca"

**Table S1** Number of unique UCE loci recovered from *in-silico* samples for 65/65 % coverage/identity percentage thresholds when matching UCE probes to contigs and number of UCE loci retained in a 50% taxon coverage matrix after trimming and filtering. Species in bold were used for probe set design

| Higher taxonomy | Species | GenBank accession No. | # UCEs (raw) | # UCEs (final) |
|---|---|---|---|---|
| Brachyura: Portunidae | *Callinectes sapidus* | GCA 020233015.1 | 1,026 | 784 |
| Brachyura: Portunidae | *Portunus trituberculatus* | GCA 017591435.1 | 1,112 | 850 |
| **Brachyura: Oregoniidae** | ***Chionoecetes opilio*** | **GCA 016584305.1** | **1,155** | **837** |
| **Brachyura: Varunidae** | ***Eriocheir sinensis*** | **GCA 013436485.1** | **1,167** | **863** |
| **Anomura: Coenobitidae** | ***Birgus latro*** | **GCA 018397915.1** | **1,196** | **840** |
| Anomura: Lithodidae | *Paralithodes camtschaticus* | GCA 018397895.1 | 949 | 682 |
| Astacidea: Cambaridae | *Procambarus clarkii* | GCA 020424385.2 | 1,156 | 828 |
| **Astacidea: Nephropidae** | ***Homarus americanus*** | **GCA 018991925.1** | **1,190** | **852** |
| Astacidea: Parastacidae | *Cherax destructor* | GCA 009830355.1 | 1,023 | 744 |
| **Achelata: Palinuridae** | ***Panulirus ornatus*** | **GCA 018397875.1** | **1,175** | **836** |
| Caridea: Atyidae | *Caridina multidentata* | GCA 002091895.1 | 919 | 680 |
| **Caridea: Palaemonidae** | ***Macrobrachium nipponense*** | **GCA 015104395.1** | **994** | **682** |
| **Dendrobranchiata: Penaeidae** | ***Penaeus japonicus*** | **GCA 017312705.2** | **1,098** | **845** |
| Isopoda: Armadillidiidae | *Armadillidium vulgare* | GCA 004104545.1 | 840 | 569 |
| Isopoda: Cirolanidae | *Bathynomus jamesi* | GCA 023014485.1 | 805 | 567 |
| Isopoda: Idoteidae | *Idotea balthica* | GCA 023373965.1 | 704 | 509 |
| Amphipoda: Hyalidae | *Parhyale hawaiensis* | GCA 001587735.2 | 851 | 556 |
| **Amphipoda: Hyalellidae** | ***Hyalella azteca*** | **GCA 000764305.4** | **1,015** | **665** |
| Amphipoda: Gammaridae | *Gammarus roeseli* | GCA 016164225.1 | 946 | 619 |
| Branchiopoda: Artemiidae | *Artemia franciscana* | GCA 019857095.1 | 381 | 143 |
| Copepoda: Temoridae | *Eurytemora affinis* | GCA 000591075.2 | 452 | 115 |

**Table S2** Assembly and UCE extraction summary statistics for the *Eumunida* dataset, including the number of assembled contigs, the number of raw UCE loci recovered from contigs, the length of the raw UCE loci, and the final number of UCE loci in a 50% taxon coverage matrix after trimming and filtering. Small letters appended to voucher numbers (*a* or *b*) denote different individuals from multi-specimen lots. Samples marked with an asterisk are included in the subset of Western Indian Ocean species, used to assess species-level variation via a “smilogram” (see Figure 2 in the main manuscript).

| Species | Voucher/ Sample ID | Year | contigs | UCEs (raw) | Locus length (bp) | | | UCEs (final) |
|---|---|---|---|---|---|---|---|---|
| | | | | | mean | min | max | |
| *Eumunida* aff. *annulosa* | MNHN-IU-2011-8607 | 2011 | 108,038 | 1,026 | 439.9 | 220 | 6,328 | 416 |
| *Eumunida annulosa* | MNHN-IU-2012-469 | 2001 | 208,899 | 962 | 409.8 | 71 | 2,840 | 383 |
| *Eumunida balssi* | USNM 150463 | 1920 | 10,556 | 272 | 282.0 | 229 | 734 | 144 |
| *Eumunida bella* | MNHN-IU-2021-311 | 1963 | 1,878 | 31 | 285.5 | 80 | 1,043 | 8 |
| *Eumunida bispinata*† | MNHN-IU-2011-5547 | 1973 | 3,014 | 28 | 256.8 | 231 | 508 | 18 |
| *Eumunida bispinata*† | MNHN-IU-2016-7024 | 2017 | 143,437 | 1,103 | 540.6 | 230 | 4,642 | 424 |
| *Eumunida capillata* | CAS:IZ:190358a | 1979 | 474,225 | 1,013 | 939.0 | 231 | 7,641 | 434 |
| *Eumunida capillata* | CAS:IZ:190358b | 1979 | 417,814 | 1,047 | 930.1 | 232 | 5,465 | 428 |
| *Eumunida capillata* | USNM 277774 | 1989 | 223 | 14 | 235.3 | 80 | 281 | 5 |
| *Eumunida capillata* | USNM 243943a | 1986 | 95 | 8 | 229.6 | 80 | 344 | 1 |
| *Eumunica capillata* | USNM 243943b | 1986 | 117 | 10 | 210.1 | 79 | 268 | 2 |
| *Eumunida capillata* | USNM 1457346 | 2017 | 334,765 | 1,048 | 1,249.3 | 231 | 6,422 | 451 |
| *Eumunida depressa* | MNHN-IU-2014-10256 | 1979 | 16,507 | 71 | 267.7 | 80 | 452 | 20 |
| *Eumunida dofleini* | USNM 1188905 | 1906 | 264 | 8 | 190.4 | 80 | 298 | 1 |
| *Eumunida funambulus* | MNHN-IU-2011-5558 | 1976 | 22,850 | 81 | 255.8 | 89 | 334 | 36 |
| *Eumunida keiji* | MNHN-IU-2011-5528 | 1985 | 5,025 | 15 | 242.1 | 83 | 332 | 6 |
| *Eumunida keiji* | MNHN-IU-20212-750b | 2020 | 25,113 | 1,078 | 895.8 | 231 | 5,011 | 443 |
| *Eumunida keiji* | MNHN-IU-20212-750a | 2020 | 152,772 | 853 | 1,278.2 | 233 | 7,116 | 401 |
| *Eumunida* aff. *keiji* | USNM 1188639 | 1972 | 199,092 | 1,082 | 783.4 | 229 | 5,385 | 434 |
| *Eumunida leavimana* | MNHN-IU-2011-5524 | 2001 | 53,206 | 905 | 357.7 | 83 | 5,023 | 396 |
| *Eumunida laevimana* | MNHN-IU-2012-310 | 1991 | 7,631 | 110 | 287.0 | 229 | 509 | 29 |
| *Eumunida macphersoni* | MNHN-IU-2014-10805 | 1979 | 2,087 | 20 | 299.8 | 80 | 538 | 2 |
| *Eumunida marginata* | MNHN-IU-2011-5446 | 2001 | 3,771 | 59 | 259.0 | 83 | 372 | 29 |
| *Eumunida minor*† | CAS:IZ:190357 | 2010 | 745,615 | 1,044 | 1,166.4 | 322 | 7,807 | 440 |
| *Eumunida minor*† | MNHN-IU-2014-19256 | 2010 | 116,423 | 1,009 | 378.8 | 219 | 5,756 | 396 |
| *Eumunida multilineata* | MNHN-IU-2011-5549 | 1984 | 1,967 | 25 | 262.7 | 83 | 461 | 7 |
| *Eumunida multispina*† | H036‡ | 2019 | 356,376 | 1,149 | 684.0 | 256 | 4,243 | 455 |
| *Eumunida multispina*† | H037‡ | 2019 | 143,500 | 796 | 1,038.8 | 234 | 4,374 | 372 |

| Species | Voucher/ Sample ID | Year | contigs | UCEs (raw) | Locus length (bp) | | | UCEs (final) |
|---|---|---|---|---|---|---|---|---|
| | | | | | mean | min | max | |
| *Eumunida pacifica* | MNHN-IU-2014-23737 | 1991 | 110,384 | 973 | 398.2 | 171 | 3,215 | 381 |
| *Eumunida cf. pacifica* | MCZ:IZ:153356 | 2019 | 510,382 | 1,043 | 1,292.4 | 81 | 12,599 | 452 |
| *Eumunida cf. pacifica* | MCZ:IZ:153353 | 2019 | 331,230 | 142 | 1,338.7 | 256 | 3,853 | 75 |
| *Eumunida picta* | MNHN-IU-2013-19053 | 2015 | 38,721 | 1,012 | 397.6 | 83 | 3,963 | 440 |
| *Eumunida* aff. *picta* | USNM 1604206 | 2017 | 123,341 | 1,108 | 1,038.4 | 240 | 4,878 | 467 |
| *Eumunida proprior* | USNM 150335 | 1909 | 16,902 | 623 | 287.4 | 219 | 502 | 286 |
| *Eumunida similior*† | MNHN-IU-2011-5516 | 1972 | 6,375 | 57 | 315.9 | 78 | 3,409 | 27 |
| *Eumunida smithii* | MNHN-IU-2014-5678 | 1991 | 2,366 | 10 | 258.0 | 83 | 430 | 2 |
| *Eumunida smithii* | MNHN-IU-2021-2751 | 2020 | 300,040 | 743 | 1,332.0 | 231 | 6,363 | 348 |
| *Eumunida spinosa* | MNHN-IU-2014-5682 | 2003 | 124,762 | 1,147 | 1,016.5 | 249 | 5,034 | 423 |
| *Eumunida spiridonovi*† | MNHN-IU-2016-8714 | 1983 | 214,685 | 77 | 1,096.9 | 242 | 8,866 | 44 |
| *Eumunida squamifera* | H001‡ | 2022 | 167,849 | 1,141 | 782.7 | 238 | 6,430 | 455 |
| *Eumunida squamifera* | MNHN-IU-2012-283 | 1984 | 17,724 | 451 | 288.2 | 229 | 1,762 | 220 |
| *Eumunida sternomaculata* | MNHN-IU-2013-14824 | 1987 | 130,497 | 996 | 481.8 | 229 | 3,808 | 382 |
| *Eumunida treguieri* | MNHN-IU-2011-5445 | 1991 | 18,337 | 308 | 302.4 | 80 | 1,956 | 161 |
| *Eumunida turbulenta* | MNHN-IU-2014-21174 | 2003 | 14,817 | 267 | 296.0 | 83 | 5,426 | 144 |
| *Eumunida* sp. | CAS:IZ:1151811 | 1998 | 5,411 | 38 | 293.2 | 80 | 1,646 | 12 |
| *Eumunida* sp. | USNM 1424077 | 2016 | 193,075 | 1,123 | 448.8 | 230 | 3,651 | 466 |
| *Eumunida* sp. | USNM 1490646 | 2018 | 6,503 | 1,094 | 278.4 | 80 | 1,061 | 458 |
| *Pseudomunida fragilis* | MCZ:IZ:151084 | 2018 | 84,402 | 957 | 1,284.1 | 229 | 6,068 | 421 |

† Sample included in the subset of Western Indian Ocean species, used to assess species-level variation via a “smilogram” (see Figure 2 in the main manuscript)
‡ Uncatalogued specimens, internal specimen ID provided

**Table S3** Assembly and UCE extraction summary statistics for the *Aratus pisonii* dataset, including sampling locality, the number of assembled contigs, the number of raw UCE loci recovered from contigs, the length of the raw UCE loci, and the final number of UCE loci in a 50% taxon coverage matrix after trimming and filtering. Lowercase letters appended to voucher numbers (*a* to *e*) denote different individuals from multi-specimen lots.

| **Locality** | **Voucher** | **Year** | **contigs** | **UCEs (raw)** | **Locus length (bp)** | | | **UCEs (final)** |
|---|---|---|---|---|---|---|---|---|
| | | | | | **mean** | **min** | **max** | |
| Hollywood, FL | MCZ:IZ:161741a | 2021 | 1,359,477 | 947 | 1,435.6 | 249 | 8,369 | 776 |
| Hollywood, FL | MCZ:IZ:161741b | 2021 | 1,190,925 | 1,007 | 1,181.5 | 308 | 6,790 | 757 |
| Hollywood, FL | MCZ:IZ:161741c | 2021 | 1,671,386 | 1,017 | 1,643.4 | 462 | 7,762 | 793 |
| Hollywood, FL | MCZ:IZ:161742b | 2021 | 1,205,609 | 971 | 1,241.6 | 334 | 8,747 | 758 |
| Hollywood, FL | MCZ:IZ:161742c | 2021 | 1,450,070 | 865 | 1,676.4 | 240 | 9,294 | 727 |
| Hollywood, FL | MCZ:IZ:161742d | 2021 | 1,241,967 | 942 | 1,318.4 | 337 | 5,837 | 757 |
| Hollywood, FL | UF 11375a | 2005 | 4,922 | 15 | 328.1 | 80 | 1,605 | 5 |
| Hollywood, FL | UF 11375b | 2005 | 26,001 | 94 | 276.6 | 80 | 497 | 50 |
| Hollywood, FL | UF 11375c | 2005 | 31,463 | 111 | 283.5 | 229 | 1,254 | 81 |
| Hollywood, FL | UF 32623a | 2005 | 94,498 | 296 | 319.3 | 229 | 3,812 | 180 |
| Miami, Virginia Key, FL | MCZ:IZ:161740a | 2021 | 1,252,535 | 917 | 1,370.4 | 311 | 12,252 | 766 |
| Miami, Virginia Key, FL | MCZ:IZ:161740b | 2021 | 1,427,479 | 897 | 1,591.4 | 392 | 6,538 | 748 |
| Miami, Virginia Key, FL | MCZ:IZ:161740c | 2021 | 1,329,040 | 960 | 1,384.4 | 347 | 6,977 | 766 |
| Miami, Virginia Key, FL | MCZ:IZ:161740d | 2021 | 1,318,555 | 918 | 1,436.3 | 392 | 7,089 | 760 |
| Miami, Virginia Key, FL | MCZ:IZ:161740e | 2021 | 1,180,088 | 994 | 1,187.2 | 310 | 10,072 | 765 |
| Miami, Virginia Key, FL | USNM 25559a | 1901 | 1,674 | 8 | 209.3 | 80 | 278 | 2 |
| Miami, Virginia Key, FL | USNM 25559b | 1901 | 17,990 | 65 | 276.8 | 80 | 801 | 35 |
| Miami, Virginia Key, FL | USNM 25559c | 1901 | 18,661 | 68 | 274.4 | 83 | 1,014 | 35 |
| Miami, Key Biscayne, FL | MCZ:IZ:161739b | 2021 | 1,517,363 | 934 | 1,624.6 | 403 | 7,093 | 765 |
| Miami, Key Biscayne, FL | MCZ:IZ:161739c | 2021 | 1,244,634 | 973 | 1,279.2 | 230 | 5,745 | 764 |
| Miami, Key Biscayne, FL | MCZ:IZ:161739d | 2021 | 1,123,724 | 954 | 1,177.9 | 269 | 6,594 | 749 |
| Miami, Key Biscayne, FL | MCZ:IZ:6194a | 1859 | 6,295 | 26 | 242.1 | 80 | 342 | 14 |
| Miami, Key Biscayne, FL | MCZ:IZ:6194b | 1859 | 5,493 | 21 | 261.6 | 80 | 340 | 9 |

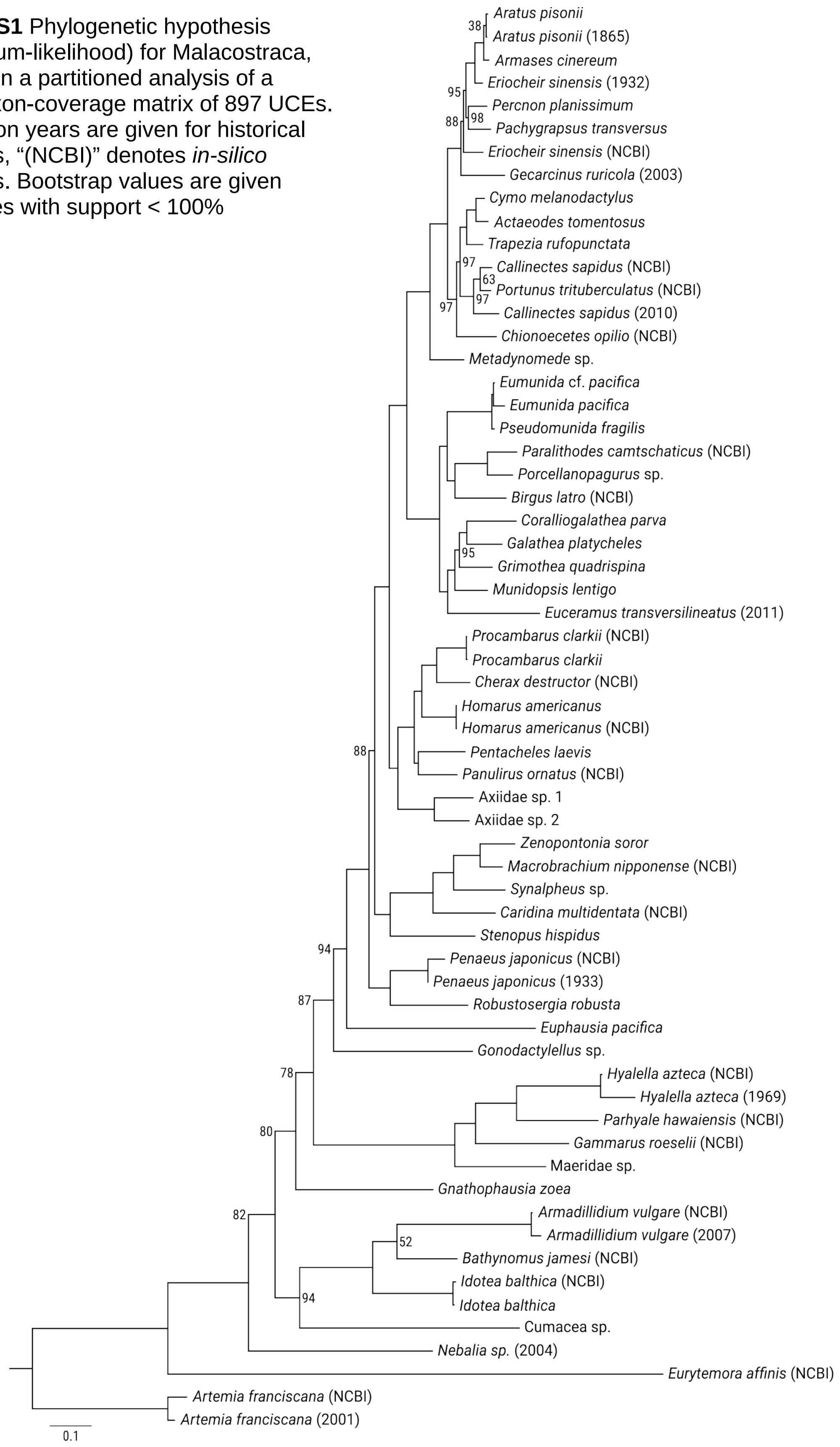


**Figure S1** Phylogenetic hypothesis (maximum-likelihood) for Malacostraca, based on a partitioned analysis of a 50% taxon-coverage matrix of 897 UCEs. Collection years are given for historical samples, "(NCBI)" denotes *in-silico* samples. Bootstrap values are given for nodes with support < 100%
